# Resting State connectivity patterns associated with trait anxiety in adolescence

**DOI:** 10.1101/2025.01.08.631901

**Authors:** Teresa Baggio, Alessandro Grecucci, Fabrice Crivello, Marc Joliot, Christophe Tzourio

**Affiliations:** Clinical and Affective Neuroscience Lab, CL.I.A.N. Lab, Department of Psychology and Cognitive Sciences (DiPSCo), University of Trento, Italy; Centre for Medical Sciences, CISMed, University of Trento, Italy; Neurofunctional Imaging Group, Institute of Neurodegenarative Disorders, UMR5293, University of Bordeaux, Bordeaux, France; Neurofunctional Imaging Group, Institute of Neurodegenarative Disorders, UMR5293, CNRS, Bordeaux, France; Neurofunctional Imaging Group, Institute of Neurodegenarative Disorders, UMR5293, CEA, Bordeaux, France; University of Bordeaux, INSERM, Bordeaux Population Health Research Center, U1219, CHU Bordeaux, Bordeaux, France; Pellegrin University Hospital Center, Bordeaux, France

**Keywords:** resting-state, adolescence, anxiety, network, default mode, connectivity

## Abstract

Anxiety symptoms can vary across different life stages, with a higher frequency during adolescence and early adulthood, increasing the risk of developing future anxiety disorders. To date, neuroscientific research on anxiety has primarily focused on adulthood, thus limiting our understanding of how anxiety may characterize earlier stages of life, and employing mostly univariate approaches, thus discounting large-scale alterations of the brain. One intriguing hypothesis is that adolescents with trait anxiety may display similar abnormalities shown by adults in brain regions ascribed to the Default Mode Network (DMN) associated with self-referential thinking, awareness, and rumination-related processes. The present study aims to expand our previous knowledge on this topic using a large sample of young individuals to uncover the resting-state connectivity patterns associated with trait anxiety in a network approach. To test our hypotheses, we analyzed the rs-fMRI images of 1263 adolescents (mean age 20.55 years) as well as their scores on anxiety trait. A significant association between trait anxiety and resting-state functional connectivity in two networks was found, with some regions overlapping with the Default Mode Network. The first network included regions such as the cingulate gyrus and the middle temporal gyri known to be involved in self-referential processing and emotional perception and control, both altered in anxiety disorders. The second network included the precuneus, possibly related to rumination that characterizes anxiety. Of note, the higher the trait anxiety, the lower the connectivity within both networks, suggesting abnormal self-referential processing, awareness, and emotion regulation abilities in adolescents with high anxiety trait. These findings provided a better understanding of the association between trait anxiety and brain rs-functional connectivity, and may pave the way for the development of potential biomarkers in adolescents with anxiety.

## Introduction

Experiencing anxiety is a common condition that almost everyone comes across during the lifetime. Nevertheless, this condition can largely vary in terms of intensity level (going from mild manifestations to clinical disorders), symptomatology (behavioural and psychological indices), frequency and occurrence. Some periods or conditions are indeed more likely to present higher anxiety level, such as early adolescence and young adulthood^1^, which are characterized by distinctive brain developmental and organizational features^2,3^.

Moreover, high levels of anxiety experienced during these periods can indeed represent a risk factor for the future development of proper anxiety disorders^4^.

In non-clinical settings anxiety is usually measured and identified through trait anxiety, a general measure of the individual propensity of perceiving stress and worry in everyday life^5^. High trait anxiety scores can point out the presence or the future development of clinical anxiety disorders^6–8^.

For this reason, the investigation of trait anxiety has been successfully employed in several studies concerning non-clinical population, analysing both structural and functional features related to it, highlighting the role of the prefrontal, the cingulate and the insular cortex, along with the amygdala^9–12^. The advantages of studying brain features associated to trait anxiety, instead of a fully diagnosed disorder, ranges from the exploration of anxiety in its subclinical forms to the exclusion of the effects of psychotropic drugs potentially taken by diagnosed patients. In this way it is possible to identify early markers of clinical disorders, studying neural predisposition factors that could impact the aetiology of anxiety.

Functional studies on trait anxiety reported impaired connectivity in brain regions associated to attentional control, such as the dorsolateral prefrontal cortex and the cingulate cortex^13,14^, and altered fronto-limbic circuits, such as impaired amygdala-prefrontal cortex connections^15^. In children and adolescents few studies have been conducted so far, finding increased activation of the right dorsolateral prefrontal cortex during a task measuring attention bias for angry faces^16^.

Moreover, a study comparing adults and adolescents’ response in threat-safety discriminations during threat reappraisal reported a decreased activity of the subgenual anterior cingulate cortex in both anxious adults and adolescents, while the ventromedial prefrontal cortex exhibited a different pattern, in particular a reduced activity in anxious adults compared to controls and an increased activity in anxious adolescents compared to controls^17^.

Despite the efforts made so far in identifying neural underpinnings of trait anxiety, most studies have employed univariate approaches including Region-of-Interest analyses, with a priori selection of specific brain regions.

However, a network approach to neuroscience may allow the identification of widespread aberrations happening in relation to a specific disorder, considering that dysfunctions in such disorders rely on distributed regions across the brain^18^.

Thus, a system neuroscience perspective can help examining how brain networks are impacted in different aberrant mental states^18,19^, investigating resting-state spontaneous fluctuations^20^. In addition, the role of brain networks in describing mental states such as anxiety is relevant also in the context of structural studies (i.e. gray and white matter) for both clinical and subclinical population^21,22^.

Anxiety in adults has been shown to influence the connectivity patterns within and between several large-scale neural networks, such as the Default Mode (DMN), the Salience (SN) and the Central Executive (CEN) (sometime also substituted by the Frontoparietal, FPN), that are considered important centres for perceptual, behavioural and emotional processing^10,23^. These three macro-networks interact with each other, in such a way that the SN, and in particular the anterior insula, is thought to mediate the switch between the CEN and the DMN, involved in externally-oriented attention the former, and internally-oriented, self-related processes the latter^18,23^.

Of these networks, the DMN seems to play a fundamental role in anxiety. Numerous studies have reported functional alterations in adults with anxiety, particularly a reduced functional connectivity in the DMN during emotion regulation tasks^24–26^. In this context, Sylvester et al.^19^ introduced a new functional network model of anxiety, proposing that anxiety behaviors and disorders result in abnormal functional connectivity pattern where increased activity in the SN is linked to decreased regulation by the DMN^24,25,27–30^. This model although interesting is limited to the observation of the brain of adults. One important question is whether the functional alterations inside the DMN may apply to adolescents with anxiety as well. The DMN is a collection of functionally related regions such as the posterior cingulate, the temporal poles, the frontal and posterior medial cortices including the precuneus.

In particular, the precuneus has been repetitively linked to anxiety^31–33^, specifically to self-referential processing rumination^32,34,35^, which consist in the persistent focus on past negative experiences and the inspection of the potential causes and consequences of a negative mood^34,36^. Moreover, also the posterior cingulate cortex has been shown to mediate anxious symptoms, being involved in emotional evaluation and having a modulatory influence on the amygdala^27,37^.

A confirmation of altered connectivity inside these regions in adolescents with trait anxiety, is crucial for advancing research in this field. Although anxiety has been a recurrent topic in the recent neuroscientific research, leading to the development of different neural hypotheses, two main limitations can be identified. First, most of the research has been conducted in small groups that are not strictly homogeneous for what concerns ages, with a net prevalence of adult subjects, making it difficult to associate anxiety features to specific sensitive periods, in particular adolescence. Since anxiety prevalence and symptomatology can vary according to the brain developmental phase, creating neural models comprising a mix of age ranges might result in an imprecise framework. Second, the general approach used to study anxiety typically involves the analysis of single, specifically localized and a priori selected brain regions, with the use of univariate methods, discounting the interpretation of cognitive and affective states through an extensive analysis of the brain’s organization. Machine learning, multivariate in nature, can help to unravel the complexity of large-scale brain features related to functional activation.

Indeed, a promising alternative to seed-based correlation concerns decomposition methods such as Independent Component Analysis (ICA), that is able to explore different resting-state networks properties for individual or group-level analyses^20^. Group ICA is a widely used strategy in resting-state analyses, given the fact that it can separate spatially or temporally independent sources, allowing for group inferences^38^, without starting from a priori spatial restraints.

Moreover, Group ICA can be useful to find potential functional network biomarkers, identifying patterns of functional activity or connectivity related to a specific condition/state^39^, thus resulting in a powerful method that in the future could be integrated with standard diagnosis approaches, which rely mostly on qualitative methods.

Given these considerations, our study aims to employ a large-scale investigation of resting-state connectivity in relation to trait anxiety in a sample of 1263 adolescents, to test the hypothesis that the Default Mode Network may be associated with trait anxiety following previous observations on adult individuals.

## Methods

### Participants

Neural data for this study was selected from the MRi-Share database^40^, a multi-modal brain MRI database acquired in a sample of 1870 young healthy adults (18-35 years old), undergoing university-level education (https://www.i-share.fr/). Informed consent was obtained from all participants and/or their legal guardians, and all the research was performed in accordance with relevant guidelines and regulations, in accordance with the Declaration of Helsinki.

The study protocol was approved by the Comite de Protection des Personnes Sud-Ouest et Outre-Mer (local ethics committee CPP SOOMIII) with agreement nr 2015-A00850-49.

The original MRi-Share database contains structural (T1 and FLAIR), diffusion (multispectral), susceptibility-weighted (SWI), and resting-state functional imaging modalities and is related to the i-Share cohort study, launched in February 2013 with the approval of the local ethics committee (CPP2015-A00850-49).

The exclusion criteria used for the MRi-Share database consists in (1) age over 35 years; (2) pregnancy or nursing; (3) claustrophobia; and (4) contraindications for head MRI, while the additional exclusion criteria used for the present study were the following: (1) diagnosis of bulimia/anorexia/obsessive compulsive disorder/depression; (2) daily alcohol consumption; and (3) drug consumption more than 10 times in life (cocaine, mdma, heroine etc..).

After the removal of subjects’ data with incidental findings requiring medical referral, and following the additional exclusion criteria for the present study, the final sample resulted to be 1263 participants (M = 341; F = 922; mean age = 20.55; SD = 2.25).

The sample characteristics are summarized in Table 1.

**Table 1.** Descriptive statistics.

|  |  |  |
| --- | --- | --- |
| **Total number** | 1263 (M = 341; F = 922) |  |
|  | **Mean** | **SD** |
| **Trait anxiety (STAI score)** | 45.08 | 10.14 |
| **Age (years)** | 20.55 | 2.25 |
| **Years of education** | 17.44 | 1.39 |

### Questionnaires

The questionnaires that participants had to fill were web-based and they were related to general physical health, lifestyle habits and mental health (see list of questionnaires at https://research.i-share.fr/data/). For the purpose of our study, the French version of the State-Trait Anxiety Inventory^5,41^ (Inventaire d’Anxiété Trait-État, Forme Y) was considered, in particular the Trait Anxiety score. The State-Trait Anxiety Inventory (STAI) is a well-validated questionnaire that measures the level of *state* and *trait* anxiety with 20 items per each, rated on a 4-point Likert scale, going from 1 “Almost Never” to 4 “Almost Always”.

Moreover, it has internal consistency coefficients ranging from .86 to .95 and test-retest reliability coefficients ranging from .65 to .75 over a 2-month interval^41^.

### MRI acquisition

Structural and functional data was acquired with a 3T Siemens Prisma scanner with a 64- channels head coil (gradients: 80 mT/m–200 T/m/sec). T1-weighted images were acquired in a time duration of 4 minutes and 54 seconds, with a voxel size of 1.0 × 1.0 × 1.0 mm^3^ (192 × 256 × 256) and key parameters 3D MPRAGE, sagittal, *R* = 2, TR/TE/TI = 2000/2.0/880 msec, while resting state data was acquired in a time duration of 14 minutes and 58 seconds, with a voxel size of 2.4 × 2.4 × 2.4 mm^3^ (88 × 88 × 66) and key parameters 2D axial, EPI, MB = 6, TR/TE = 850/35.0 msec, flip angle = 56°, fat sat^40^. Before the rs-fMRI acquisition, participants were asked to “keep their eyes closed, to relax, to refrain from moving, to stay awake, and to let their thoughts come and go”^40^.

### Pre-processing

Resting state fMRI data was pre-processed through CONN Toolbox^42^ (RRID:SCR_009550) release 21.a. Functional and anatomical data were preprocessed using a flexible preprocessing pipeline^43^ including realignment with correction of susceptibility distortion interactions, slice timing correction, outlier detection, direct segmentation and MNI-space normalization, and smoothing. Functional data were realigned using SPM realign & unwarp procedure^44^, where all scans were coregistered to a reference image, and resampled using b-spline interpolation to correct for motion and magnetic susceptibility interactions. Potential outlier scans were identified using ART^45^, and a reference BOLD image was computed for each subject by averaging all scans excluding outliers. Functional and anatomical data were normalized into standard MNI space, segmented into grey matter, white matter, and CSF tissue classes, and resampled to 2 mm isotropic voxels following a direct normalization procedure^46,47^. Last, functional data were smoothed using spatial convolution with a Gaussian kernel of 4 mm full width half maximum (FWHM)^48^.

In addition, functional data were denoised using a standard denoising pipeline^43^ including the regression of potential confounding effects characterized by CSF, motion parameters and their first order derivatives, outlier scans, white matter timeseries, and linear trends within each functional run, followed by bandpass frequency filtering of the BOLD timeseries^49^ between 0.008 Hz and 0.09 Hz.

### Group ICA analysis

In the first-level analysis, Group ICA (independent component analysis) estimated 20 temporally coherent networks from the rs-fMRI data combined across all subjects. This number was chosen according to previous studies and CONN default suggestion. The resulting networks were then visually inspected for noise control^50^.

Group ICA analysis is able to compute a series of spatial maps characterizing the BOLD response across a set of independent spatial components, using Calhoun’s group-level ICA approach^38^, with variance normalization pre-conditioning, optional subject-level dimensionality reduction, subject/condition concatenation of BOLD signal data along temporal dimension, group-level dimensionality reduction (to the target number of dimensions/components), fastICA for estimation of independent spatial components, and GICA3 backprojection for individual subject-level spatial map estimation^43^ (See Figure 1 for a visual representation of the Group ICA steps).

**Figure 1.**
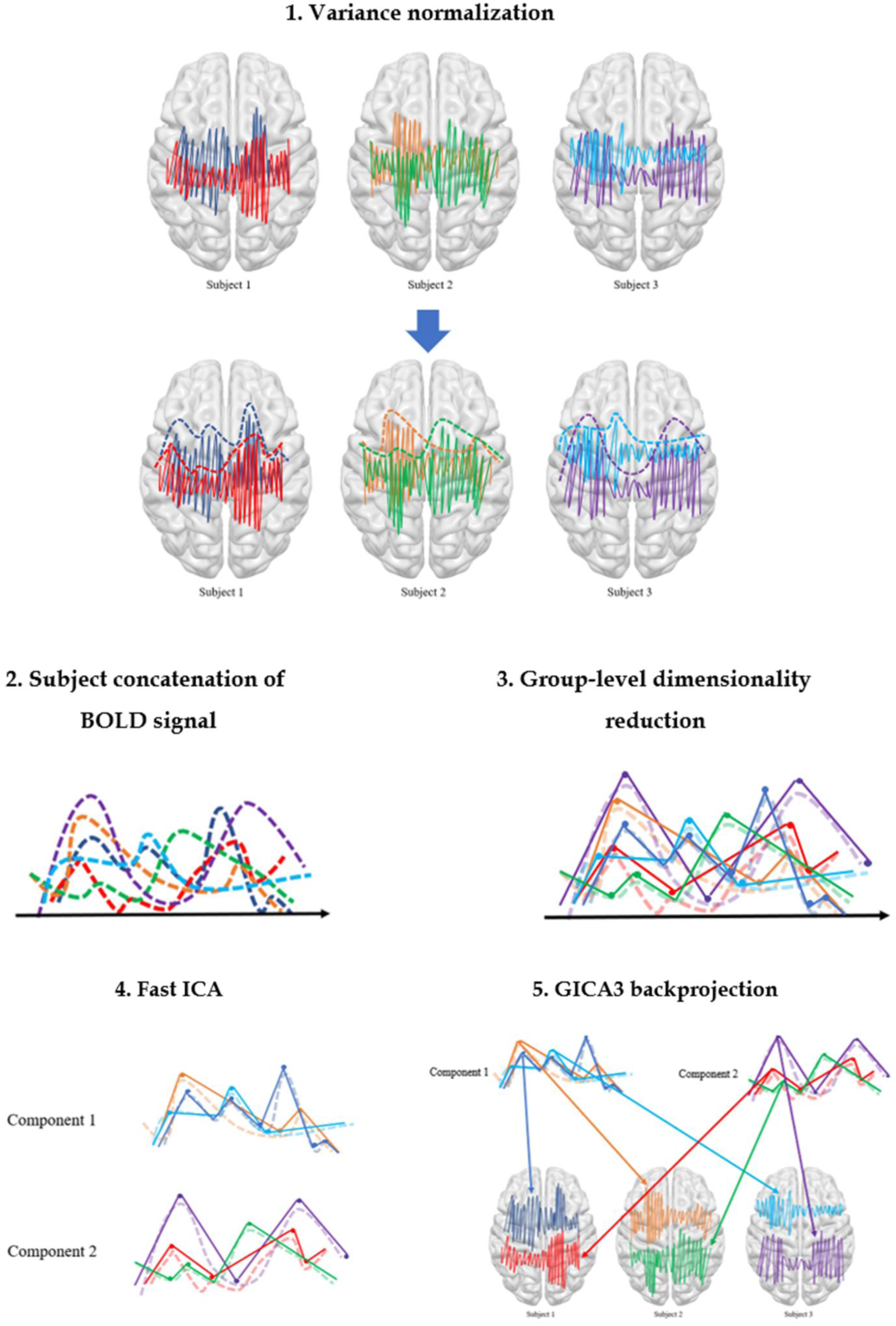
Group ICA steps *Group ICA steps automatically performed by CONN include: 1) variance normalization to adjust the variance of the time series, aiming at rendering data more comparable across subjects; optional subject-level dimensionality reduction (here not represented); 2) subject/condition concatenation of BOLD signal data along temporal dimension, in order to combine together BOLD signal data of different subjects/conditions along the time axis; 3) group-level dimensionality reduction, which refers to the reduction of the data complexity in order to extract common features or patterns; 4) fastICA for the estimation of the independent components from the mixed data matrix; 5) GICA3 backprojection, consisting in the estimation of individual subject-level spatial map starting from group-level spatial map. For the purpose of the image, here only three subjects and two components are represented*.

The components thus represent an estimation of independent hemodynamic sources, and have a specific contribution for each voxel signal^38^. In the second-level analysis, for each component a GLM contrast was performed in order to calculate how trait anxiety influences resting state connectivity, controlling for sex. The contrast was set with a voxel level p <0.001 uncorrected and a cluster level p < 0.05 pFDR corrected. Significant results were plotted in Surf Ice (https://www.nitrc.org/projects/surfice/) and BrainNet Viewer^51^ for visualization.

The anatomical specification of the significant areas was derived from MRIcroGL Toolbox^52^, with the integrated AAL atlas function, and CONN automatic atlas (Harvard-Oxford atlas).

## Results

### Group ICA analysis

Of the twenty estimated components, the connectivity between two networks, e.g. IC 8 and IC 10, and specific brain regions resulted to be significantly associated with trait anxiety (See figure 2 and 4 for the visual representation). In particular, the higher the trait anxiety, the lower the connectivity of these two networks with specific brain regions (See Figure 3 and 5, Table 2 and 3 for the visual and anatomical specification of the significant areas resulting from the 2^nd^ level contrast). IC 8 included the superior temporal gyrus, the inferior frontal gyrus, the middle occipital gyrus, the precuneus, the postcentral gyrus, the middle temporal gyrus, the fusiform gyrus, and the hippocampus. This network showed a decreased resting-state connectivity with the posterior cingulate gyrus, and the middle temporal gyrus. IC 10 included the cerebellum (vermis), the inferior and middle occipital gyrus, the precuneus, the cuneus, the calcarine sulcus, and the lingual gyrus. This network showed a decreased resting-state connectivity with the precuneus.

**Figure 2.**
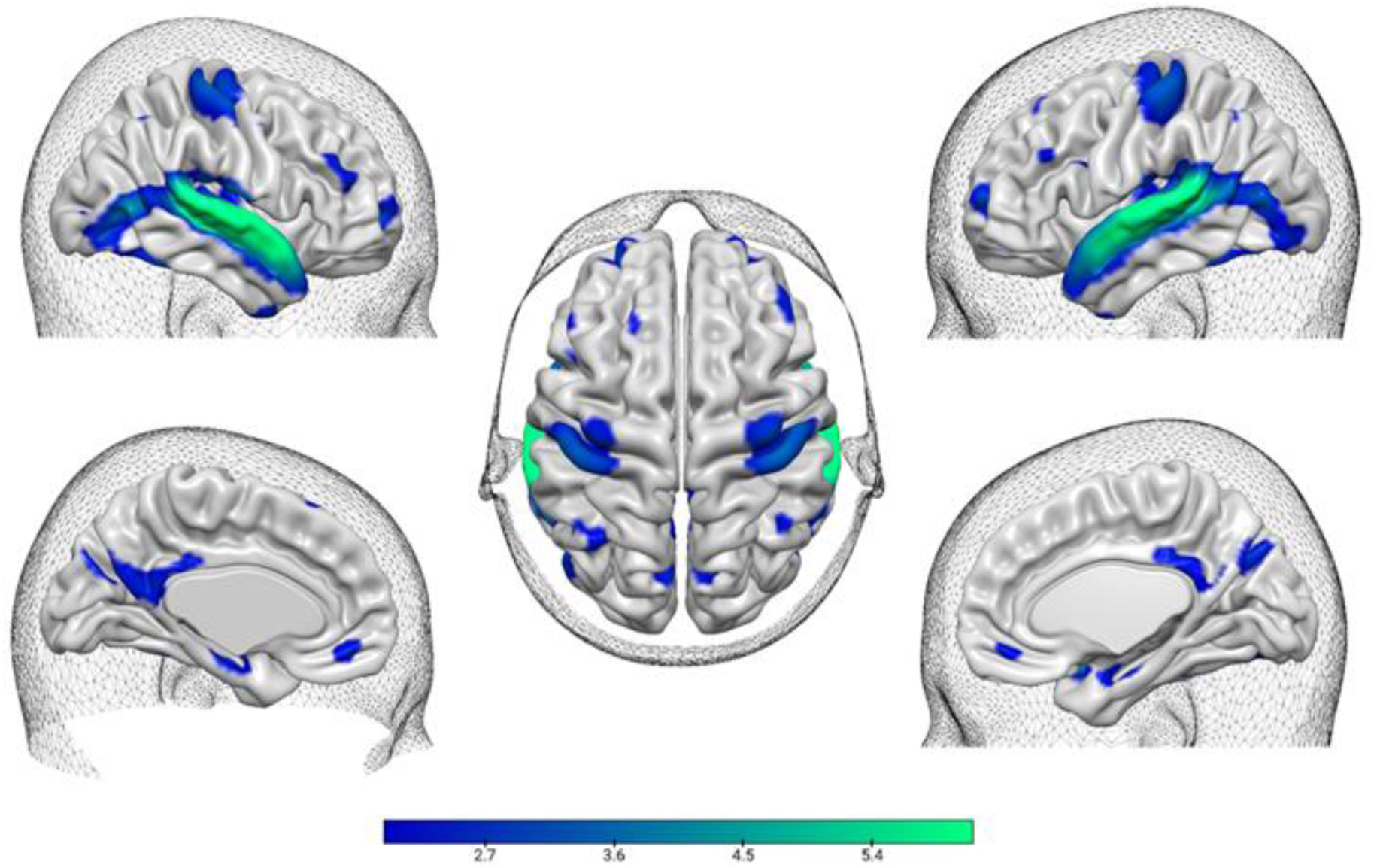
Visual representation of the component of interest IC 8 *Components IC 8. The colours refer to the z score derived from the statistical values obtained by the analysis, for each brain area*.

**Figure 3.**
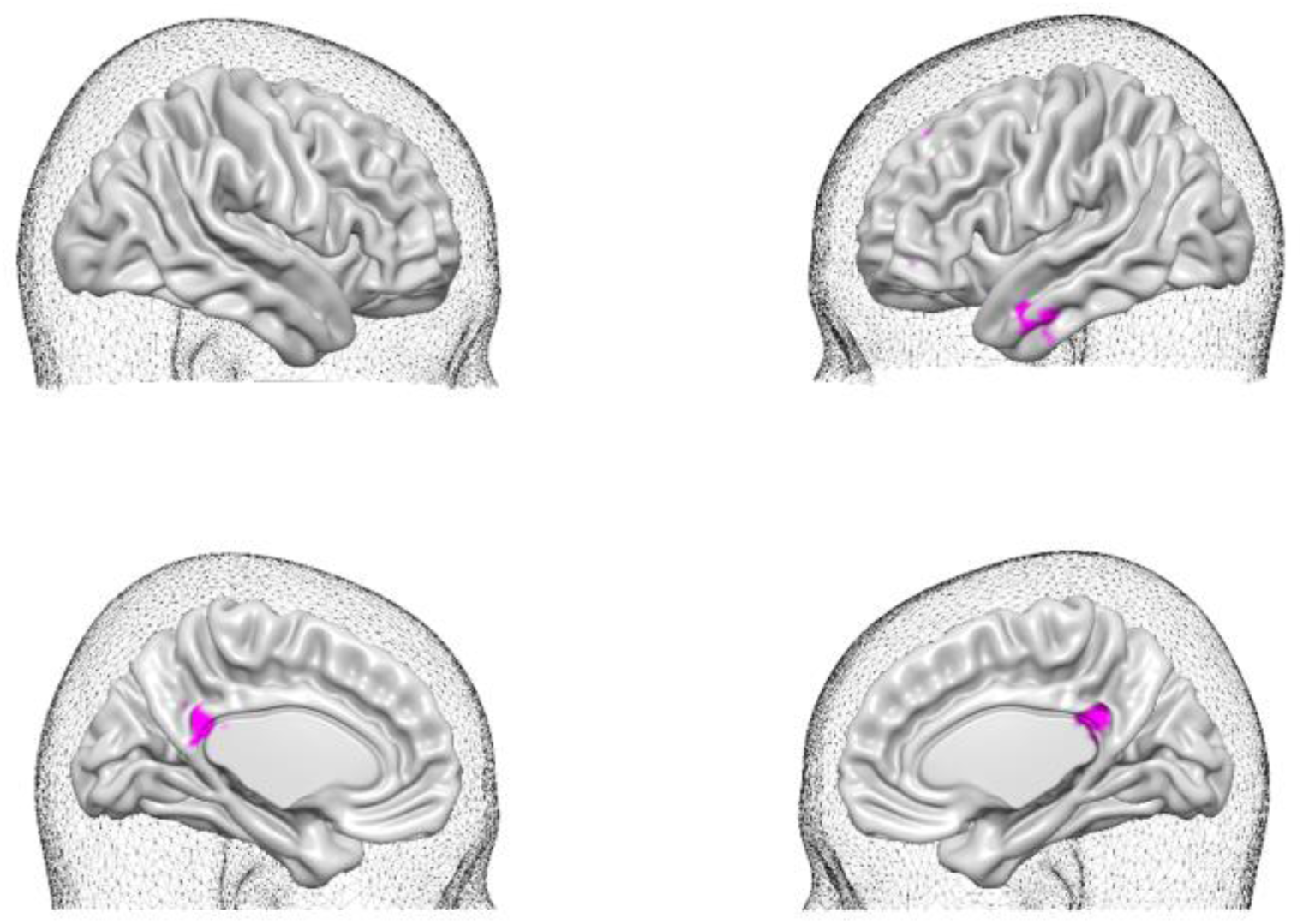
Visual representation of the areas showing decreased resting-state connectivity with IC8 *Significant areas resulted from the 2^nd^ level contrast. The coloured regions represent areas showing a decreased resting-state connectivity with IC8, associated with increase trait anxiety*.

**Figure 4.**
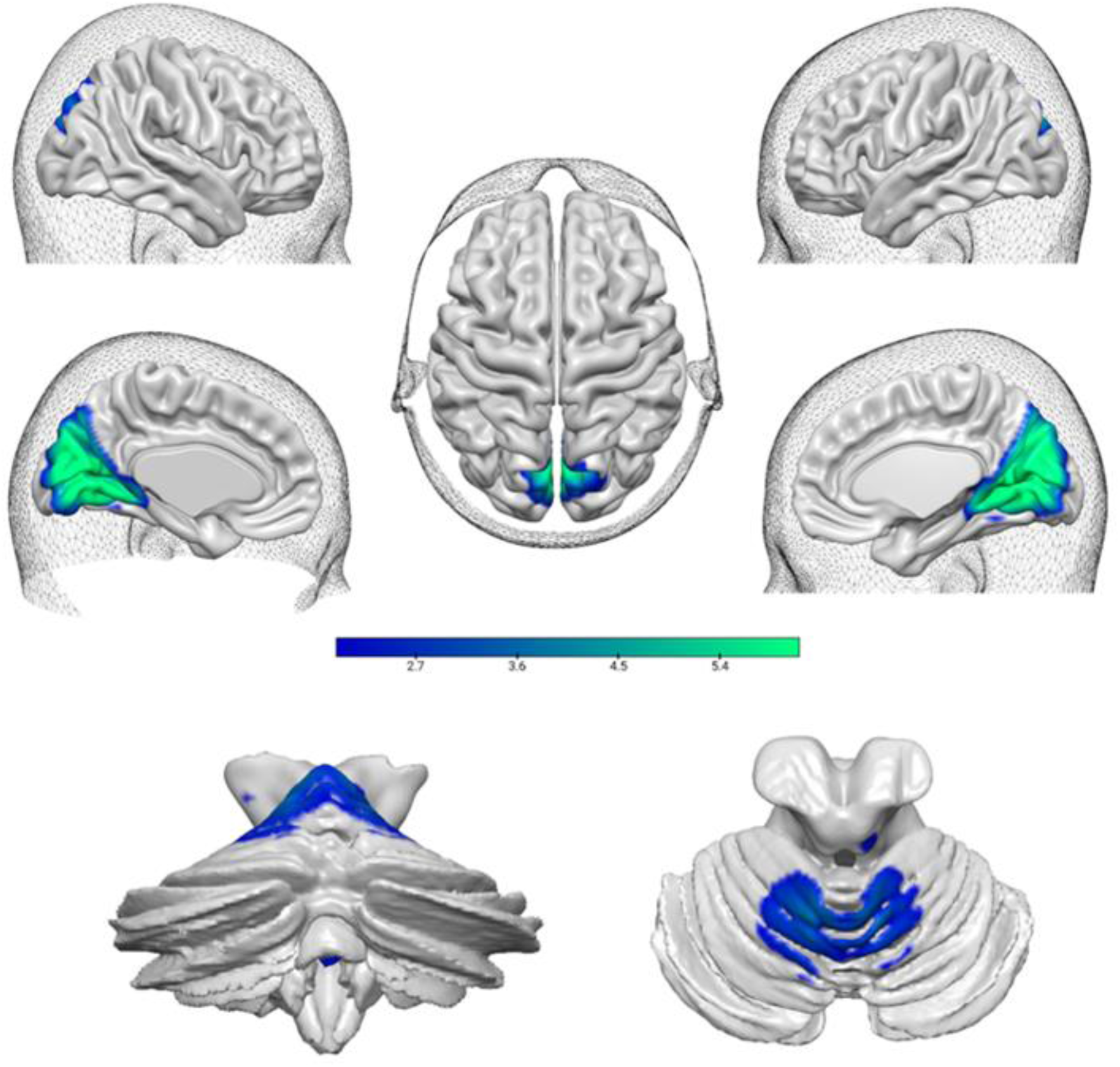
Visual representation of the component of interest IC 10 *Components IC 10. The colours refer to the z score derived from the statistical values obtained by the analysis, for each brain area*. *On top, cortical and subcortical areas. At the bottom, posterior cerebellar view (left) and superior cerebellar view (right)*.

**Figure 5.**
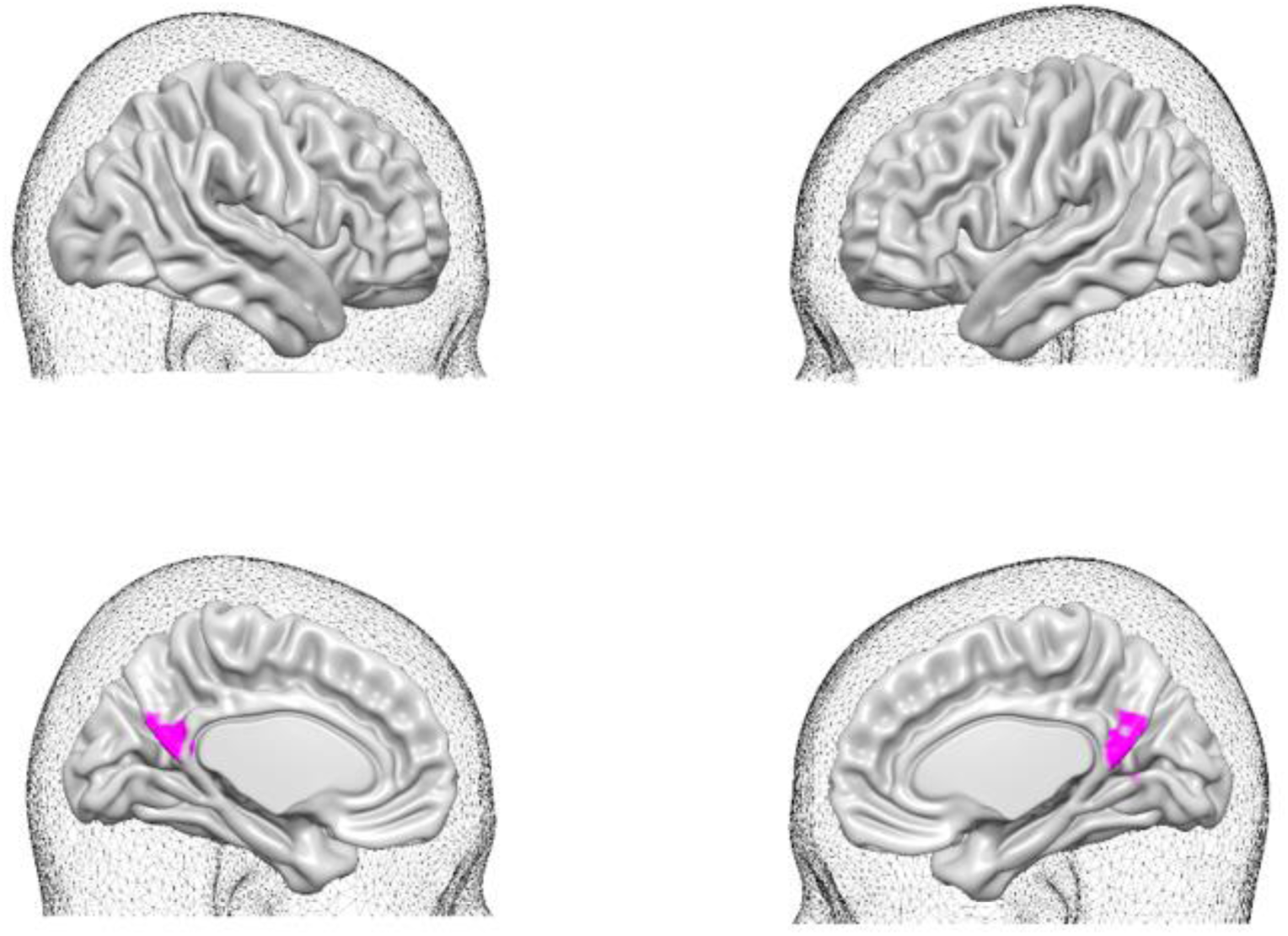
Visual representation of the areas showing decreased resting-state connectivity with IC10 *Significant areas resulted from the 2^nd^ level contrast. The coloured regions represent areas showing a decreased resting-state connectivity with IC10, associated with increase trait anxiety*.

**Table 2.** Anatomical specification of the areas showing decreased resting-state connectivity with IC8. *Significant areas resulted from the 2^nd^ level contrast. The anatomical denomination is based on Harvard-Oxford Atlas. All the above-mentioned areas showed decreased resting-state functional connectivity with the reported IC, in association with trait anxiety*.

| IC | ROI | Voxels | Peak Statistics<br>(z) | MNI coordinates<br>(x, y, z) |
| --- | --- | --- | --- | --- |
| 8 | Cingulate Gyrus, posterior division | 52 | 4.4 | (-2,-44,+24) |
|  | Middle Temporal Gyrus, anterior division, Left | 56 | 4.2 | (-58,-4,-26) |

**Table 3.** Anatomical specification of the areas showing decreased resting-state connectivity with IC10.

| IC | ROI | Voxels | Peak Statistics<br>(z) | MNI coordinates<br>(x, y, z) |
| --- | --- | --- | --- | --- |
| 10 | Precuneous | 149 | 4.3 | (+2,-58,+18) |

*Significant areas resulted from the 2<sup>nd</sup> level contrast. The anatomical denomination is based on Harvard-Oxford Atlas. All the above-mentioned areas showed decreased resting-state functional connectivity with the reported IC, in association with trait anxiety.*

## Discussion

Our study aimed to explore the relationship between trait anxiety and resting-state functional connectivity in a large cohort of young participants, investigating this relationship in association with the Default Mode Network, using Group ICA. The analysis revealed that two independent components, IC8 and IC10, showed significant associations between trait anxiety and decreased connectivity in several key brain regions.

In the following sections we will explain in detail such results.

### A temporo-parietal network

IC8 encompassed regions associated with higher order cognitive and sensory processes that are integral to several networks and showed a decreased resting state connectivity with regions ascribable to the posterior cingulate gyrus and the middle temporal gyrus.

Indeed some of the regions comprising the network, such as the precuneus, the superior, and middle temporal gyrus, are regions of the Default Mode Network, involved in social cognition and working memory, and previous research have shown a decreased functional connectivity of them in highly anxious individuals, also at rest^10^. Moreover, an alteration of the DMN has been shown to be a common feature of anxiety disorders^10,53,54^, in particular a reduced connectivity of it during rest has been associated to deficits in self-evaluation and introspection^10^. The postcentral gyrus contributes to somatosensory processing^10^, whereas the hippocampus has been suggested to have a role also in contextual anxiety and memory, being strongly connected with other brain regions involved in emotion regulation, including the amygdala, the prefrontal cortex and the hypothalamus^55–57^. Together, the regions included in IC8 suggest a network that might be involved in the integration of sensory information with cognitive processes. Notably, this network showed decreased connectivity in the middle temporal gyrus, as well as the posterior cingulate gyrus. These regions are also integral to the DMN, implicated in self-referential thinking and emotional regulation.

The middle temporal gyrus is indeed involved in mind wandering, autobiographical memory retrieval, semantic memory and social cognition^58,59^; moreover, the left middle temporal gyrus have been found impaired in young patients with social anxiety disorders^60^.

The posterior cingulate cortex has been associated with internally directed cognition, self-referential processes, focus of attention regulation and environmental monitoring for threatening stimuli^61–63^, being a key region that interacts also with different neural networks, such as the fronto-parietal, the sensorimotor and the dorsal attention networks^64^. Additionally, abnormalities in functional connectivity of the posterior cingulate cortex have been documented in several psychiatric and neurological conditions, including attention deficit hyperactive disorder, autism spectrum disorder and schizophrenia^64^.

Therefore, we hypothesize that a decreased resting-state connectivity of IC8 network with the middle temporal gyrus and posterior cingulate gyrus might be related to disrupted self-referential processes such as self-focus, and maladaptive rumination, often observed in anxiety disorders, as both the posterior cingulate cortex and the precuneus are involved in ruminative processes^34,65^.

### An occipito-cerebellar network

On the other side, IC10 encompassed regions associated with sensory integration, emotion regulation, and visual processing including the cerebellar vermis, inferior and middle occipital gyrus, precuneus, cuneus, calcarine sulcus, and lingual gyrus. The cerebellum, beyond its well-known role in motor coordination, plays a key role in various cognitive, affective and social functions^66^, and has been increasingly linked to anxiety-related processes^67^. Specifically, the cerebellar vermis, often referred to as the “limbic cerebellum”, is critical for emotional regulation due to its strong connections with key limbic system regions, such as periaqueductal gray and dorsal raphe nuclei, both of which are associated with anxiety modulation and stress responses^67^. Moreover, cerebellar structure was associated with social abilities in a sample of children and teenagers, providing support for the social and cognitive role of cerebellum^68^. Regions within the occipital lobe, such as inferior and middle occipital gyri, cuneus, and calcarine sulcus, are primarily involved in visual perception and processing of sensory stimuli, particularly under conditions of uncertainty^69,70^. Disruptions in functional connectivity within these areas have been associated with anxiety disorders, possibly due to impaired sensory integration and heightened sensitivity to uncertain stimuli, as demonstrated by a decreased functional connectivity in visual processing areas such as the lingual and the inferior occipital gyrus in patients with social anxiety disorders, suggesting perceptual impairments^71^. Overall, this network, involved in visuo-sensory and affective processes, showed decreased connectivity in the precuneus, the key default mode region involved in self-processing, consciousness and episodic memory retrieval^72^.

The posterior precuneus has many structural pathways connected to the visual and dorsal attention networks, suggesting its role as a connector among multiple large-scale networks^73^. Studies in patients with social anxiety disorder have revealed decreased connectivity between the left precuneus and some areas such as posterior lobe of cerebellum, inferior temporal gyrus, parahippocampal gyrus and medial prefrontal cortex^31^. These disruptions may contribute to the social and emotional processing difficulties seen in such disorder. Similarly, panic disorder patients showed altered regional homogeneity in the precuneus^32^. In addition, high trait anxious individuals displayed decreased activation in the precuneus, during anticipation of uncertain threat compared to the certain condition^74^.

We thus hypothesize that a decreased resting-state connectivity of IC10 network with the precuneus might be involved in disruptions in the ability to process internal thoughts, self-awareness and visual perceptual processing, as trait anxiety could lead to cognitive perturbations in areas related to visual-sensory integration and emotion regulation.

## Conclusions and limitations

In conclusion, this study provided new evidence of the association between trait anxiety and decreased resting-state functional connectivity in two key independent networks, including areas ascribable to the default Mode Network, using a large sample of adolescents. Our findings underscore the importance of large-scale brain networks in understanding the neural correlates of anxiety and point toward disruptions in self-referential and sensory integration processes as key mechanisms underlying anxiety-related cognitive dysfunctions. From a methodological point of view, this study aligns with the recent trend in system neuroscience to study affective and pathological states at a network level^18^. To do this we took advantage of an unsupervised machine learning algorithm known as Group ICA, instead of the classically used methods such as ROI to ROI or seed-based connectivity. This method has the advantage of being multivariate in nature and does not require an a priori selection of brain areas. As such it allows greater sensitivity to detect distributed network-level functional changes.

However, some limitations must be acknowledged. First, the findings should be extended with caution to proper anxiety disorders, as the current study focuses on trait anxiety rather than clinical diagnoses. Additionally, controlling for other demographic variables, such as socioeconomic status, would ensure that the results are not confounded by these factors.

Addressing these limitations will enhance the generalizability and robustness of our findings in understanding the neural underpinnings of anxiety in adolescents.

## Author contributions

T.B.: conceptualization, methodology, formal analysis, writing—original draft. A.G.: conceptualization, methodology, writing—original draft, supervision. F.C.: data acquisition, writing—original draft, supervision. M.J.: data acquisition, writing—original draft, supervision. C.T.: data acquisition, supervision. All authors have read and agreed to the published version of the manuscript.

## Data availability statement

Due to French regulations regarding sharing of the medical imaging data, individual raw data used for this study cannot be shared through a public repository. Rather, to have access to i-Share and MRi-Share de-identified data, please contact Christophe Tzourio. A request has to be submitted to the i-Share Scientific Collaborations Coordinator with a letter of intent (explaining the rationale and objectives of the research proposal), and a brief summary of the planned means and options for funding. The i-Share Steering Committee will assess this request, and provide a response (principle agreement, request to reformulate the application or for further information, refusal with reasons). If positive, applicants will have to complete and return an application package that will be reviewed by the principal investigator, the Steering Committee, and the operational staff. Reviews are based on criteria such as the regulatory framework and adherence to regulations (access to data, confidentiality), the scientific and methodological quality of the project, the relevance of the project in relation to the overall consistency of the cohort in the long term, the complementarity/competition with projects planned or currently underway, ethical aspects. Both de-identified raw and processed data (and data dictionaries) will be shared after (i) final approval of the application, and (ii) formalization of the specifics of the collaboration.

